# OmicsFUSION: A Pretrained Hyena-Based Framework for Encoding, Reconstruction, and Representation of the Omics and DNA Data

**DOI:** 10.1101/2025.09.05.674329

**Authors:** Nazar Beknazarov, Nikita Pavlichenko, Artem Bashkatov, Maria Poptsova, Alan Herbert

## Abstract

Recent advances in Natural Language Processing (NLP) have spurred the application of Large Language Models (LLMs) to bioinformatics, enabling innovative approaches to DNA sequence encoding. However, genomic function is not solely determined by the primary DNA sequence—it also depends on complex, multi-layered omics data. This auxiliary data is inherently sparse, structured across multiple tracks, and poses a challenge for traditional unimodal approaches. Simultaneously, many bioinformatics tasks demand a unified signal source that encapsulates this information, rather than requiring researchers to input each omics feature individually. To overcome these limitations, we introduce a novel framework for the integrated representation of DNA sequences and their associated omics data. Central to our approach is a large-scale, batch-structured omics dataset optimized for deep learning at scale. Our framework is built around four key novelties: (1) a new model architecture that jointly processes DNA and omics signals; (2) an extension of Masked Language Modeling (MLM) to omics tracks for effective data reconstruction; (3) single-nucleotide embeddings that fuse all input modalities; and (4) interval-level embeddings that summarize broader genomic regions. We release a collection of pretrained models capable of reconstructing, embedding, and generating unified representations of DNA enriched with functional omics context.

**Code:** https://github.com/aaai-2025-submission/anomymous_submission_aaai2025

**Model and Datasets:** https://drive.google.com/drive/folders/10hcuq4rTCr8bKQFqkpz7R91MGtKvR0Va?usp=sharing

## Introduction

Recent advancements in NLP, exemplified by models such as GPT-3 and LLaMA, have catalyzed parallel developments in bioinformatics (Devlin et al. 2019; Brown et al. 2020; Touvron et al. 2023). A significant area of need within this field is the development of a unified model or data vector that encapsulates comprehensive genomic information, which can then be applied to solve various downstream tasks. Pioneering pre-trained LLMs like DNABERT and ESM have significantly impacted the field, giving rise to new models and methodologies that rely on the embeddings or representations generated by these pre-trained systems (Ji et al. 2021; Rives et al. 2021; Lin et al. 2023). Such approaches have been particularly effective for DNA sequences, whose linear structure closely resembles that of natural language texts. However, unlike in NLP, DNA encoders face a unique challenge: genomic information in a living cell is not solely encoded by the primary DNA sequence but is also heavily influenced by a range of complex omics features.

Recent progress in DNA encoders has largely focused on adapting the MLM approach, where DNA sequence regions are masked, and the model is trained to predict the missing nucleotides. This has proven to be effective for capturing the syntactic and semantic information inherent to the DNA sequence itself. However, these models neglect the rich functional context provided by other layers of genomic encoding. Although DNA LLMs excel at understanding sequence level, they cannot resolve the ambiguity where identical DNA sequences exhibit different functional roles due to epigenetic modifications or other regulatory signals.

This limitation becomes particularly apparent when considering tissue specificity. Since the DNA sequence is identical across nearly all cell types within an organism, sequence-only models are fundamentally incapable of distinguishing between different tissues or cell lineages. Consequently, building tissue-specific predictive models requires a different approach. In contrast, omics features vary significantly between tissues, providing the molecular basis for their distinct identities and functions. Therefore, a model that incorporates omics data is not only better suited for general genomic tasks but is essential for the development of accurate tissue-specific models (Buenrostro et al. 2013; Klemm, Shipony, and Greenleaf 2019).

Omics features, particularly epigenetic markup, represent a diverse set of signals that are crucial to understanding genome functioning. These features include DNA methylation, various histone modifications, RNA polymerase binding sites, chromatin accessibility, and transcription factor and chromatin binding sites, among others (Cedar and Bergman 2009; Bannister and Kouzarides 2011; Buenrostro et al. 2013). This data is characterized by its high dimensionality, sparsity, and organization into multiple, often discontinuous, tracks along the genome. It is well-established that these signals contain unique information essential for regulating cellular processes. Alterations in these omics landscapes can profoundly influence cell behavior and are associated with numerous diseases. Although the DNA sequence provides the foundational template for these marks, an additional layer of genomic information is encoded independently at the epigenome level. Numerous attempts to predict the full spectrum of omics features from the DNA sequence alone have had limited success. Even recent largescale foundation models fail to achieve satisfactory performance, underscoring the need to directly incorporate omics data into these models. This gap has motivated us to develop a model that extends the advancements in DNA encoding while incorporating omics features as a primary input.

The development of such a model presents a powerful opportunity to streamline genomic research. This enables researchers requiring epigenomic context to use a single comprehensive embedding instead of manually integrating numerous individual data tracks. Furthermore, predictive models trained on this unified representation could enable novel in silico research, such as studying the functional impact of single-nucleotide variants on cellular behavior. This capability has profound implications for mutation research, personalized medicine, and the fundamental decoding of previously unknown regulatory dependencies within the genome.

To realize this vision, we addressed several fundamental challenges. First, we needed a large-scale, curated dataset of diverse omics experiments. For this work, we compiled approximately 40,000 individual experimental tracks for the mouse genome and 60,000 for the human genome, grouped into nearly 500 and 1,000 distinct categories, respectively. Efficiently managing this extensive dataset - both on disk and in memory for GPU-based training - presents significant engineering challenges, especially when handling the data gaps characteristic of experimental omics datasets. Moreover, adapting MLM to omics data required developing a novel masking strategy that accounts for both the multi-track structure and inherent sparsity of omics datasets. A new architecture was also necessary to effectively process these inputs. Additionally, creating robust embeddings for both single nucleotides and genomic intervals demanded further research into how best to integrate these disparate data types into a unified representation.

To overcome these challenges, we developed and validated a novel framework for learning a joint, multimodal representation of DNA sequences and their corresponding omics profiles. Our approach uses a specialized architecture with an extended masked language modeling objective to co-embed these data types, producing unified genomic representations that are more predictive and biologically meaningful than sequence-only embeddings.

### Novelty and Contributions

The novelty of this work is multifaceted and can be summarized as follows:

1. **New Datasets**: We introduce two new, large-scale omics datasets for mouse and human genomes, batched and optimized for deep learning pipelines, enabling multi-GPU and multi-node training.
2. **Integrated Framework**: We provide a Lightning and PyTorch-based framework that facilitates the training of new models for real-world tasks using our developed architectures.
3. **Novel Architecture**: We have designed a new, efficient architecture specifically for processing multimodal omics data.
4. **Masking Design**: A new masking strategy has been developed for MLM-based tasks tailored to the unique characteristics of omics data.
5. **Pre-trained Models**: We offer a range of pre-trained encoder and embedding models with varying latent space dimensions (378, 512, and 768) to suit different computational and task-specific needs.
6. **Omics Track Prediction**: Our models have been used to predict and test omics data that were not included in the training set, demonstrating their generalizability.

## Related Work

### Language Models for Biological Sequences

The application of NLP models to bioinformatics has been driven by the powerful analogy of the “genome as language.” This paradigm treats DNA sequences as linear constructs similar to text, enabling the use of architectures like the Transformer. Foundational models such as DNABERT adapted the MLM technique, where regions of the DNA sequence are masked and the model learns to predict the missing nucleotides, thereby capturing the “syntax” of the genome without a need for labeled data (Devlin et al. 2019; Ji et al. 2021; Rogers, Kovaleva, and Rumshisky 2020). This approach proved to be effective for creating embeddings applicable to various downstream tasks. However, these initial models faced challenges with the unique structure of genomic data, such as the inefficiency of k-mer tokenization and the computational expense of the self-attention mechanism, which scales quadratically with the sequence length, limiting the ability to model long-range dependencies. These limitations, combined with concerns about overparameterization, sometimes resulted in underperformance compared to simpler baseline models (Compeau, Pevzner, and Tesler 2011; Du et al. 2019).

Subsequent architectural innovations sought to overcome these limitations. The development of HyenaDNA replaced the costly attention mechanism with long implicit convolutions, enabling the model to process context windows of up to one million tokens at single-nucleotide resolution (Nguyen et al. 2023). This leap in efficiency and granularity is crucial for genomics, where functional elements can be separated by large distances and a single nucleotide change can have significant consequences. While these advancements have pushed the boundaries of what sequence-only models can achieve, they still encounter a fundamental ceiling. The core issue is that genomic function is not determined by the DNA sequence alone. Identical DNA sequences can play different functional roles in different tissues due to epigenetic regulations - a critical biological context that sequence-only models inherently fail to capture. This inherent ambiguity necessitates the integration of additional data layers for complete biological understanding.

Later, pre-trained LLMs in bioinformatics were trying to fill this gap by incorporating SNP data or by increasing the number of genomes from different species. The most notable examples here are Nucleotide Transformer (Dalla-Torre et al. 2025), which was trained on 3,202 human genomes and 850 genomes from diverse species, and Evo2 (Brixi et al. 2025), which was trained on 16,704 eukaryotic genomes, all representative prokaryotic genomes, 41,253 metagenomes and metagenome-assembled genomes, and 33,457 organelle genomes.

### Multimodal Learning in Computational Biology

The limitations of unimodal models have catalyzed a shift towards multimodal learning, which aims to create more holistic understanding by integrating diverse data types (Huang et al. 2022; Hu et al. 2023; Zhou et al. 2024). Biological systems are inherently multimodal, with function arising from a complex interplay between the genome, epigenome, transcriptome, and proteome. Fusing these different data “views” allows models to leverage complementary information, reinforce signals, and, most importantly, uncover complex interactions that are invisible to any single modality. Deep learning has become the primary tool for this task due to its ability to learn non-linear relationships across high-dimensional and heterogeneous data sources. The development of such integrated models is not just an incremental improvement, but a necessary step to overcome the performance plateaus faced by even the most advanced sequence-only approaches.

### Representation learning

The task of generating representations for a range of tokens, as opposed to individual tokens, has long posed a significant challenge to researchers. A pioneering development in NLP was the BERT framework, which introduced a special classification token, [CLS], added to the input sequence (Vaswani et al. 2017; Devlin et al. 2019). The embedding of this token [CLS], originally intended for sequence-level classification tasks such as next-sentence prediction, was subsequently adopted as a de facto embedding of sentences. While the original paper suggested that the [CLS] token’s pre-training objective improved overall model quality, later research did not consistently find evidence for its superiority as a sentence representation vector.

Consequently, alternative strategies emerged that moved away from relying on a single designated token. These approaches focused on working with the output embeddings of all tokens in a sequence directly. Various pooling strategies were explored, with studies finding that a simple mean pooling of all token embeddings can serve as a robust substitute for the [CLS] embedding, often demonstrating comparable or even superior performance. Such methods also offered the flexibility to generate representations for text spans that extend beyond natural sentence delimiters, an approach well suited for bioinformatics. The genome, for instance, contains numerous functional elements with overlapping and nested relationships, making a priori segmentation a difficult task.

In this context, contrastive learning has garnered considerable attention for training robust representations. For example, as shown in SimCSE(Gao, Yao, and Chen 2021), using dropout as a source of noise to create positive pairs can be an effective training objective . While this specific technique showed limited success in our particular task, the underlying principle of contrastive learning proved highly effective. Frameworks such as PSC-CPI have successfully applied contrastive objectives to protein fragments of varying lengths, maximizing the similarity between different views of the same protein (a positive pair) against views from other proteins (negative pairs) (Wu et al. 2024). Similarly, our approach utilizes a contrastive loss on positive pairs of embeddings that represent different context lengths, while treating other instances within the same batch as negative pairs. This strategy aligns with the latest state-of-the-art representation learning models, such as E5 and BGE-M3, which emphasize the critical importance of dataset quality and the careful selection of meaningful positive pairs over architectural or pooling innovations, suggesting that this is a key factor for achieving superior performance (Wang et al. 2022; Multi-Granularity 2024).

### Model benchmarking

The surge in the number of different LLMs in bioinformatics has made it difficult for researchers to compare them properly. Different objectives, training datasets, and architectures have generated models that predict different outcomes. This trend raised the need for standardized tasks and datasets on which these models can be compared. In this work, we measured the model performance on benchmark datasets published in the following work (Grešová et al. 2023):

- Human Enhancers Cohn (`human_enhancers_cohn`): Adapted from a 2018 study by Cohn et al., this dataset contains 27,791 human enhancer sequences, which are DNA regions that regulate gene activity. The sequences have a median length of 500 base pairs.
- Human Enhancers Ensembl (`human_enhancers_ensembl`): This dataset was built using 154,842 human enhancer sequences from the FANTOM5 project via the Ensembl database. Its negative samples were randomly generated from the human genome.
- Human Nontata Promoters (`human_nontata_promoters`): From a 2017 paper by Umarov and Solovyev, this dataset includes 36,131 non-TATA promoter sequences of a fixed 251 base-pair length. Negative samples are random fragments from human gene bodies.
- Human OCR Ensembl (`human_ocr_ensembl`): Contains 174,756 sequences representing Open Chromatin Regions (OCRs) from the Ensembl Regulatory Build. These are accessible DNA regions identified by DNase-seq experiments. Negative samples were randomly generated from the human genome.

## Methods

### Problem Statement

The overall model architecture is presented in Figure 1. The model was specifically designed to address the following objectives:

**Figure 1:**
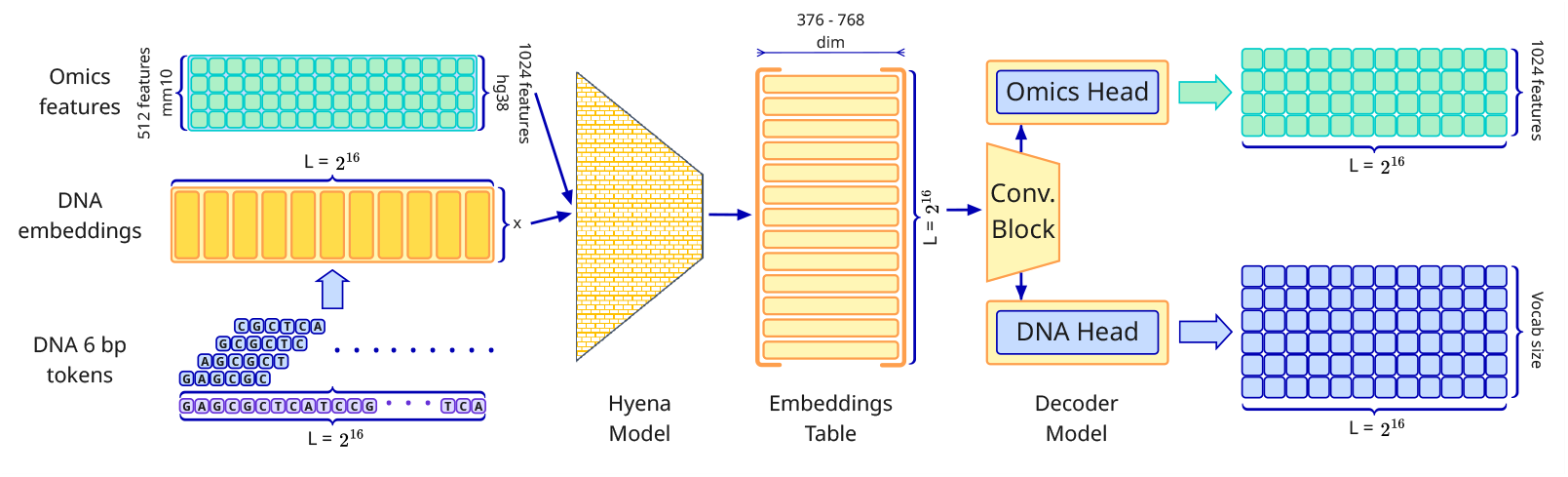
A general model overview

- Encode both DNA sequences and omics features into a unified vector representation that captures the full breadth of the input data.
- Ensure that the encoder captures a broad genomic context, allowing for long-range dependencies to be modeled.
- Design a decoder capable of reconstructing the omics data with an F1 score exceeding 80%.
- Enable the generation of embeddings not only for single base pairs but also for broader genomic regions.

### DNA Encoding

This study utilizes two genome assemblies: `hg38` for human and `mm10` for mouse. A standard DNA encoding scheme was applied. The DNA sequences were divided into overlapping k-mers of length 6, with each 6-mer treated as an individual token, following the methodology described in (Ji et al. 2021).

### Omics Encoding

One of the primary sources of omics data was ChipAtlas (Zou, Ohta, and Oki 2024). From this resource, we collected over 60,000 independent experiments for the human genome and more than 40,000 for the mouse genome. Each track in this database corresponds to a specific omics feature, such as histone modifications, RNA polymerase binding, transcription factor occupancy, and chromatin accessibility. Each experiment was performed on a specific tissue type or a specific cell line. Due to data availability constraints, this study focused exclusively on tissue-specific experiments.

Each omics track was obtained in BED format, with signal values ranging from 0 to 1000, reflecting the peak intensity of detected signals across genomic intervals. To standardize the input, all values were normalized to the [0, 1] range by dividing by 1000. Aggregated omics signals are visualized in Figure 2. Aggregation was performed to mitigate artifacts from individual experiments and to produce more robust, representative input tracks.

**Figure 2:**
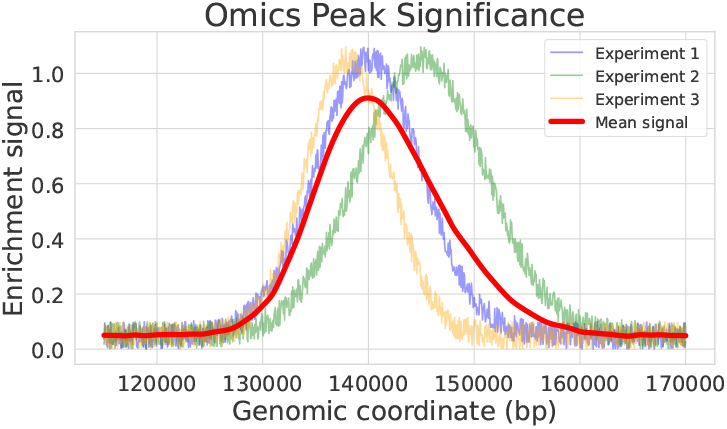
Omics data signal

**Figure 3:**
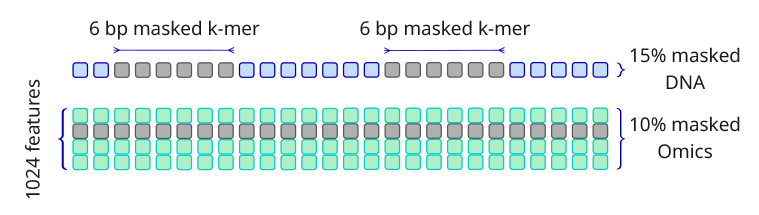
Masking pattern.

For preprocessing, data were grouped by an omics feature and tissue type. Groups with fewer than three distinct experiments were excluded from the training set. For the remaining groups, we aggregated the signals by taking the maximum value across all experiments within each group. This procedure yielded a single representative BED file per group, which served as the input for downstream processing. The final dataset included over 1,000 aggregated tracks for the human genome and approximately 500 for the mouse genome.

### Data Chunking

To improve data access efficiency, entire chromosomes were divided into fixed-size intervals of length *W*, where *W* = 2^16^ in this work. Each interval was intersected with the BED tracks using a method analogous to `bedtools intersect` . The resulting intervals, along with their corresponding signal values and source track identifiers, were stored in a new plain text format. This format was chosen for its simplicity and ease of testing.

During both training and evaluation, only the data corresponding to the currently processed interval is loaded into memory, significantly reducing data transfer overhead. Additionally, we excluded all blacklisted regions (Amemiya, Kundaje, and Boyle 2019).

### Train-Test Split

To prevent data leakage and overfitting, the train-test split was performed independently of the data content. The entire genome was evenly partitioned into short subregions of length *w*. The model was then trained incrementally with increasing context lengths, starting from *w* = 2^8^ and progressing to *w* = 2^16^. Due to the large number of subregions, effective stratification was achieved without the need for additional heuristics.

### Masking and Missing Data Handling

A significant proportion of omics features are not uniformly available across all tissues. These missing features are encoded identically to the masked ones by replacing them with zeros.

For DNA, masking was performed following the BERT strategy, where 15% of the tokens are masked. Due to our use of overlapping tokens, no individual token was masked in isolation; instead, contiguous spans of at least 6 tokens were masked to preserve the structural consistency of the k-mer encoding.

Standard masking strategies are not directly applicable to omics data. This is because neighboring omics signal values tend to be highly correlated, making random token-level masking ineffective. To address this, we adopted an alternative strategy: a random Boolean vector was generated, where no more than 10% of its entries were set to zero. This vector was used to mask entire omics signals, replacing selected features with zeros across the entire region.

### Model Architecture

The overall matrix dimensions and data flow are illustrated in Figure 1.

#### DNA Encoder Block

The DNA encoder follows a standard design. Tokens are embedded via an embedding layer, and rotary positional encodings (RoPE) (Su et al. 2024) are applied to incorporate positional context. The resulting representations are projected into the desired embedding dimensionality using a fully connected layer.

#### Omics Encoder Block

The omics features are projected directly into the same embedding dimensionality as the DNA tokens using a single fully connected layer.

#### Encoder Architecture

The encoder consists of *K*_encoder_ layers composed of Hyena blocks. In the final configuration of the model, 12 layers were used.

The Hyena block is a substitute for the attention mechanism. It replaces the standard 𝒪 (*L*^2^) time and space complexity of attention with 𝒪 (*L* log *L*) time and 𝒪 (*L*) space complexity for similar input dimensions (Poli et al. 2023). This allows for a substantial increase in context length under constrained computational budgets. While the Hyena architecture has seen limited adoption in the NLP domain, it has demonstrated broader applicability in bioinformatics (Nguyen et al. 2023; Dalla-Torre et al. 2025). For this reason, we also benchmarked the Hyena block against attention-based alternatives.

#### Decoder Architecture

The decoder consists of a shared initial component followed by two separate heads: one for omics reconstruction and one for DNA reconstruction. Unlike the encoder, which operates on a broad context window, the decoder is restricted to a narrower receptive field.

The shared component is implemented as a stack of three convolutional layers. Multiple architectures for the two heads were evaluated, each responsible for mapping the vectors to the predictions. Among all configurations tested, the ConvNeXt block (Woo et al. 2023) achieved the best performance and was selected as the final design.

The overall encoder-decoder structure can be summarized as:

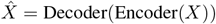

where *X* ∈ ℝ ^*L×d*^ denotes the ground truth input matrix, and 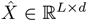 represents the model’s predictions. Here, *L* is the sequence length and *d* is the number of omics features.

#### Loss Selection

For DNA reconstruction, we employed the standard *Cross Entropy Loss* applied to masked tokens. For omics data, two types of loss functions were explored: *Mean Squared Error (MSE) Loss* and *Binary Cross Entropy (BCE) Loss*. Early experiments demonstrated that *BCE Loss* yielded superior performance, showing better ability to reconstruct values within the [0, 1] interval.

To emphasize the importance of masked targets during training, we applied a weighting scheme: masked omics targets were given three times the weight of unmasked targets.

The overall loss functions are defined as follows:

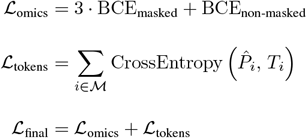

Here, 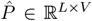 represents the probability distribution of the predicted token at position *i*, while *T* ∈ ℕ*^L^*denotes the true token index at that position. *V* is the vocabulary size. *ℳ* - set of indices of masked tokens.

### Representation Learning

The representation learning model was trained independently of the main model. The encoder component was inherited from the main model and kept frozen during training.

The embeddings generated by the encoder were directly fed into a separate representation embedding model. Each training batch consisted of two sets of genomic intervals - designated as left and right. The following conditions were enforced for each batch:

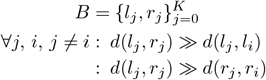

Here, *B* represents a batch containing *K* entity pairs, where *K* is determined by GPU memory constraints. Each *l*_*i*_ and *r*_*i*_ denotes a genomic interval, defined by five parameters: chromosome number, start index, end index, strand, and tissue. The similarity metric *d* was implemented as a dot product in this work.

The left and right intervals were generated using two distinct strategies:

- **Proximity-Based Intervals**. Left and right intervals were generated as follows:
  1. The parent intervals were sampled uniformly throughout the genome from random tissues and strands.
  2. Intervals overlapping blacklisted regions were excluded.
  3. Two subregions were generated within each parent interval.
  4. Subregions included a variety of types: nested, adjacent, intersecting, and parallel (from opposite strands).
  5. One subregion was assigned as the left interval, the other as the right.
- **Function-Based Intervals**. To enable embeddings that capture genomic function, intervals were generated based on omics features:
  1. A random omics feature was selected.
  2. Two independent genomic regions containing the selected feature were sampled.
  3. Both regions were cropped to the current window size.
  4. One region was designated as the left interval, the other as the right.

#### Region Embedding

Region embeddings are computed via simple mean pooling, applied independently to the embeddings of the left and right intervals. This approach ensures that each region is represented by a fixed-size vector that captures the average signal across its span.

#### Representation Loss

The contrastive loss used for training the representation model is defined as follows:

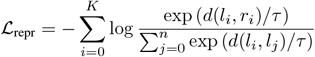

Here, *d*(*l*_*i*_, *r*_*i*_) denotes the similarity between the left and right embeddings of the *i*-th pair, computed using the dot product. The temperature parameter *τ* controls the sharpness of the distribution. The denominator sums over all left-region embeddings in the batch, treating them as negative samples relative to *l*_*i*_, while the numerator encourages high similarity between the paired intervals (*l*_*i*_, *r*_*i*_).

## Experimental Results

### Dataset

The primary source of omics data for this study was the ChipAtlas (Zou, Ohta, and Oki 2024) repository. We provide the full pipeline for downloading, aggregating, and processing the raw ChipAtlas data. The final aggregated dataset used in this publication is also publicly released. For the DNA component, we utilized the hg38 and mm10 genome assemblies, which are publicly available and did not require additional preprocessing or aggregation.

### Training Details

#### Hardware and Software

All training was conducted on a Slurm-managed high-performance computing cluster comprising a mix of hardware configurations. This included nodes equipped with H100 GPUs (1 GPU per node) and A100 GPUs (8 GPUs per node). The number of CPU cores per node varied; in practice, each GPU was allocated 6 CPU cores. Due to fluctuating cluster availability, the number of active nodes and GPUs varied dynamically throughout the training process. In total, over 50,000 GPU hours were consumed across all experiments.

The training pipeline was implemented using `PyTorch 2.3` and orchestrated via `Lightning 2.4` . Data loading was optimized using `torchdata` streaming datasets. All training was performed using BF16 mixed precision to accelerate computation and reduce memory overhead.

#### Pre-training of the Encoder–Decoder

We trained six primary encoder–decoder models corresponding to the configurations presented in Table 1. Model training followed a staged curriculum that progressively increased the input context length:

**Table 1:** Model size by embedding dimension and genome dataset

| Genome | Emb. Size | Total Param. | Size (MB) |
| --- | --- | --- | --- |
| hg38 | 768 | 99.1M | 396 |
| mm10 | 768 | 98.3M | 393 |
| hg38 | 512 | 47.0M | 188 |
| mm10 | 512 | 46.5M | 185 |
| hg38 | 384 | 28.5M | 113 |
| mm10 | 384 | 29.1M | 112 |

| Stage | Context $L$ | Epoch num |
| --- | --- | --- |
| 1 | $2^9$ | 2 |
| 2 | $2^{13}$ | 2 |
| 3 | $2^{16}$ | 2 |

Optimization was performed using the **AdamW** optimizer with hyperparameters *β*_1_ = 0.9 and *β*_2_ = 0.95. We employed a peak learning rate of 10^−3^, linear warm-up for 25

#### Representation Model

For representation learning, the encoder weights were frozen, and a 5-layer ConvNeXt architecture (hidden size 768) was trained using contrastive learning. Training was conducted on 100,000 randomly sampled genomic intervals using the `AdamW` optimizer with a learning rate of 1 × 10^−3^ and a temperature parameter *τ* = 1. Each batch consisted of *K* = 128 positive interval pairs, while negative samples were drawn implicitly from the remaining batch elements.

#### Metrics

For DNA token reconstruction, accuracy served as the primary evaluation metric. For omics data reconstruction, we reported accuracy, F1 score, and ROC AUC. In the representation learning setting, we measured quality using in-batch recall among the top-*N* most similar elements.

#### Results

Table 3 presents the core training metrics. As expected, the performance on non-masked tokens and omics features surpasses that on their masked counterparts. Nevertheless, the model achieves strong performance even on the masked sets, indicating robust generalization to unseen genomic regions and omics tracks.

Table 2 compares OmicsFUSION against benchmark models across several standard tasks. Our framework consistently outperforms existing baselines, demonstrating its effectiveness in integrating DNA and omics data.

**Table 2:** Performance comparison between benchmark models and OmicsFUSION

| Dataset | Benchmark |  | OmicsFUSION |  |
| --- | --- | --- | --- | --- |
|  | Accuracy (%) | F1 Score (%) | Accuracy | F1 Score |
| human_enhancers_cohn | 68.9 | 71.3 | 84.4 | 84.3 |
| human_enhancers_ensembl | 81.1 | 74.6 | 86.4 | 81.2 |
| human_nontata_promoters | 86.5 | 84.4 | 88.6 | 88.8 |
| human_ocr_ensembl | 68.8 | 72.0 | 86.8 | 78.9 |

**Table 3:**
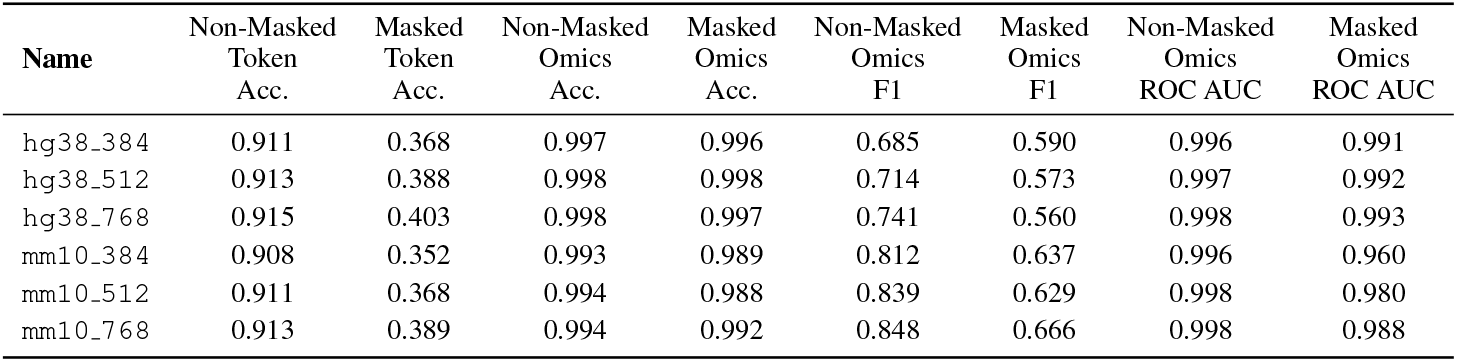
Evaluation metrics for different models on hg38 and mm10 datasets. Accuracy, F1, and ROC AUC are reported for both masked and non-masked omics regions reconstruction and DNA token reconstruction tasks. All metrics are calculated on test set.

#### Representation Quality

We evaluated two representation learning strategies and found that the omics-based approach yielded superior results, highlighting the importance of functional context in generating informative embeddings.

#### Runtime Efficiency

Thanks to a range of optimizations, the training pipeline executed reliably across varying GPU and node configurations. The use of Hyena blocks in place of traditional attention mechanisms enabled efficient distributed training using the Distributed Data Parallel strategy, even on GPUs with 80GB of memory.

## Conclusion and Future Work

OmicsFUSION surpasses existing sequence-only language models on external genomics benchmarks, while maintaining computational efficiency. These results support the central premise of this work: jointly modeling DNA sequences and epigenomic signals produces more informative and generalizable genomic representations.

Future work will focus on analyzing the model’s predictions of unseen omics tracks and extending the framework to a broader set of downstream tasks. Additionally, we aim to incorporate interpretability methods to gain deeper insight into model behavior and improve biological understanding.

## Notes

### Competing Interest Statement

The authors have declared no competing interest.

## References

Amemiya, H. M.; Kundaje, A.; and Boyle, A. P. 2019. The ENCODE blacklist: identification of problematic regions of the genome. Scientific reports, 9(1): 9354.

Bannister, A. J.; and Kouzarides, T. 2011. Regulation of chromatin by histone modifications. Cell research, 21(3): 381–395.

Brixi, G.; Durrant, M. G.; Ku, J.; Poli, M.; Brockman, G.; Chang, D.; Gonzalez, G. A.; King, S. H.; Li, D. B.; Merchant, A. T.; et al. 2025. Genome modeling and design across all domains of life with Evo 2. BioRxiv, 2025–02.

Brown, T.; Mann, B.; Ryder, N.; Subbiah, M.; Kaplan, J. D.; Dhariwal, P.; Neelakantan, A.; Shyam, P.; Sastry, G.; Askell, A.; et al. 2020. Language models are few-shot learners. Advances in neural information processing systems, 33: 1877–1901.

Buenrostro, J. D.; Giresi, P. G.; Zaba, L. C.; Chang, H. Y.; and Greenleaf, W. J. 2013. Transposition of native chromatin for fast and sensitive epigenomic profiling of open chromatin, DNA-binding proteins and nucleosome position. Nature methods, 10(12): 1213–1218.

Cedar, H.; and Bergman, Y. 2009. Linking DNA methylation and histone modification: patterns and paradigms. Nature Reviews Genetics, 10(5): 295–304.

Compeau, P. E.; Pevzner, P. A.; and Tesler, G. 2011. How to apply de Bruijn graphs to genome assembly. Nature biotechnology, 29(11): 987–991.

Dalla-Torre, H.; Gonzalez, L.; Mendoza-Revilla, J.; Lopez Carranza, N.; Grzywaczewski, A. H.; Oteri, F.; Dallago, C.; Trop, E.; de Almeida, B. P.; Sirelkhatim, H.; et al. 2025. Nucleotide Transformer: building and evaluating robust foundation models for human genomics. Nature Methods, 22(2): 287–297.

Devlin, J.; Chang, M.-W.; Lee, K.; and Toutanova, K. 2019. Bert: Pre-training of deep bidirectional transformers for language understanding. In Proceedings of the 2019 conference of the North American chapter of the association for computational linguistics: human language technologies, volume 1 (long and short papers), 4171–4186.

Du, J.; Jia, P.; Dai, Y.; Tao, C.; Zhao, Z.; and Zhi, D. 2019. Gene2vec: distributed representation of genes based on coexpression. BMC genomics, 20(Suppl 1): 82.

Gao, T.; Yao, X.; and Chen, D. 2021. Simcse: Simple contrastive learning of sentence embeddings. arXiv preprint arXiv:2104.08821.

Grešová, K.; Martinek, V.; Čechák, D.; Šimeček, P.; and Alexiou, P. 2023. Genomic benchmarks: a collection of datasets for genomic sequence classification. BMC Genomic Data, 24(1): 25.

Hu, F.; Hu, Y.; Zhang, W.; Huang, H.; Pan, Y.; and Yin, P. 2023. A multimodal protein representation framework for quantifying transferability across biochemical downstream tasks. Advanced Science, 10(22): 2301223.

Huang, K.; Fu, T.; Gao, W.; Zhao, Y.; Roohani, Y.; Leskovec, J.; Coley, C. W.; Xiao, C.; Sun, J.; and Zitnik, M. 2022. Artificial intelligence foundation for therapeutic science. Nature chemical biology, 18(10): 1033–1036.

Ji, Y.; Zhou, Z.; Liu, H.; and Davuluri, R. V. 2021. DNABERT: pre-trained Bidirectional Encoder Representations from Transformers model for DNA-language in genome. Bioinformatics, 37(15): 2112–2120.

Klemm, S. L.; Shipony, Z.; and Greenleaf, W. J. 2019. Chromatin accessibility and the regulatory epigenome. Nature Reviews Genetics, 20(4): 207–220.

Lin, Z.; Akin, H.; Rao, R.; Hie, B.; Zhu, Z.; Lu, W.; Smetanin, N.; Verkuil, R.; Kabeli, O.; Shmueli, Y.; et al. 2023. Evolutionary-scale prediction of atomic-level protein structure with a language model. Science, 379(6637): 1123–1130.

Multi-Granularity, M.-L. M.-F. 2024. M3-Embedding: Multi-Linguality, Multi-Functionality, Multi-Granularity Text Embeddings Through Self-Knowledge Distillation.

Nguyen, E.; Poli, M.; Faizi, M.; Thomas, A.; Wornow, M.; Birch-Sykes, C.; Massaroli, S.; Patel, A.; Rabideau, C.; Bengio, Y.; et al. 2023. Hyenadna: Long-range genomic sequence modeling at single nucleotide resolution. Advances in neural information processing systems, 36: 43177–43201.

Poli, M.; Massaroli, S.; Nguyen, E.; Fu, D. Y.; Dao, T.; Baccus, S.; Bengio, Y.; Ermon, S.; and Ré, C. 2023. Hyena hierarchy: Towards larger convolutional language models. In International Conference on Machine Learning, 28043–28078. PMLR.

Rives, A.; Meier, J.; Sercu, T.; Goyal, S.; Lin, Z.; Liu, J.; Guo, D.; Ott, M.; Zitnick, C. L.; Ma, J.; et al. 2021. Biological structure and function emerge from scaling unsupervised learning to 250 million protein sequences. Proceedings of the National Academy of Sciences, 118(15): e2016239118.

Rogers, A.; Kovaleva, O.; and Rumshisky, A. 2020. A Primer in BERTology: What we know about how BERT works. arXiv. Preprint posted online on February, 27.

Su, J.; Ahmed, M.; Lu, Y.; Pan, S.; Bo, W.; and Liu, Y. 2024. Roformer: Enhanced transformer with rotary position embedding. Neurocomputing, 568: 127063.

Touvron, H.; Lavril, T.; Izacard, G.; Martinet, X.; Lachaux, M.-A.; Lacroix, T.; Roziere, B.; Goyal, N.; Hambro, E.; Azhar, F.; et al. 2023. Llama: Open and efficient foundation language models. arXiv preprint arXiv:2302.13971.

Vaswani, A.; Shazeer, N.; Parmar, N.; Uszkoreit, J.; Jones, L.; Gomez, A. N.; Kaiser, Ł.; and Polosukhin, I. 2017. Attention is all you need. Advances in neural information processing systems, 30.

Wang, L.; Yang, N.; Huang, X.; Jiao, B.; Yang, L.; Jiang, D.; Majumder, R.; and Wei, F. 2022. Text embeddings by weakly-supervised contrastive pre-training. arXiv preprint arXiv:2212.03533.

Woo, S.; Debnath, S.; Hu, R.; Chen, X.; Liu, Z.; Kweon, I. S.; and Xie, S. 2023. Convnext v2: Co-designing and scaling convnets with masked autoencoders. In Proceedings of the IEEE/CVF conference on computer vision and pattern recognition, 16133–16142.

Wu, L.; Huang, Y.; Tan, C.; Gao, Z.; Hu, B.; Lin, H.; Liu, Z.; and Li, S. Z. 2024. Psc-cpi: Multi-scale protein sequence-structure contrasting for efficient and generalizable compound-protein interaction prediction. In Proceedings of the AAAI conference on artificial intelligence, volume 38, 310–319.

Zhou, Y.; Geng, P.; Zhang, S.; Xiao, F.; Cai, G.; Chen, L.; Initiative, A. D. N.; and Lu, Q. 2024. Multimodal functional deep learning for multiomics data. Briefings in Bioinformatics, 25(5): bbae448.

Zou, Z.; Ohta, T.; and Oki, S. 2024. ChIP-Atlas 3.0: a datamining suite to explore chromosome architecture together with large-scale regulome data. Nucleic Acids Research, 52(W1): W45–W53.

